# Denitrification by bradyrhizobia under feast and famine and the role of the bc1 complex in securing electrons for N_2_O reduction

**DOI:** 10.1101/2022.09.29.510233

**Authors:** Yuan Gao, Magnus Øverlie Arntzen, Morten Kjos, Lars R. Bakken, Åsa Frostegård

## Abstract

Rhizobia living as microsymbionts inside nodules have stable access to carbon substrates, but also have to survive as free-living bacteria in soil where they are starved for carbon and energy most of the time. Many rhizobia can denitrify, thus switch to anaerobic respiration under low O_2_ tension using N-oxides as electron acceptors. The cellular machinery regulating this transition is relatively well-known from studies under optimal laboratory conditions, while little is known about this regulation in starved organisms. It is, for example, not known if the strong preference for N_2_O-over NO_3_ ^-^-reduction in bradyrhizobia is retained under carbon limitation. Here we show that starved cultures of a *Bradyrhizobium* strain with respiration rates 1-18% of well-fed cultures, reduced all available N_2_O before touching provided NO_3_^-^. Proteomics showed similar abundance of Nap (periplasmic NO_3_ ^-^ reductase) and NosZ (N_2_O reductase), suggesting that competition between electron pathways to Nap and NosZ favoured N_2_O reduction also in starved cells, similar to well-fed cultures. This contrasts the general notion that NosZ activity diminishes under carbon limitation. The results suggest that bradyrhizobia carrying NosZ can act as strong sinks for N_2_O under natural conditions and that this criterion should be considered in the development of biofertilizers.

**Importance:** Legume cropped farmlands account for substantial N_2_O emissions globally. Legumes are commonly inoculated with N_2_-fixing bacteria, rhizobia, to improve crop yields. Rhizobia belonging to *Bradyrhizobium*, the micro-symbionts of several economically important legumes, are generally capable of denitrification but many lack genes encoding N_2_O reductase and will be N_2_O sources. Bradyrhizobia with complete denitrification will instead act as sinks since N_2_O-reduction efficiently competes for electrons over nitrate reduction in these organisms. This phenomenon has only been demonstrated under optimal conditions and it is not known how carbon substrate limitation, which is the common situation in most soils, affects the denitrification phenotype. Here we demonstrate that bradyrhizobia retain their strong preference for N_2_O under carbon starvation. The findings add basic knowledge about mechanisms controlling denitrification and support the potential for developing novel methods for greenhouse gas mitigation based on legume inoculants with the dual capacity to optimize N_2_-fixation and minimize N_2_O emission.

## Introduction

Bacteria in most natural and engineered environments are faced with fluctuating availability of nutrients and need to adapt to a “feast and famine” lifestyle. While many soil types are rich in total organic carbon, the concentration of bioavailable carbon substrate is low, particularly in non-rhizosphere soil where lack of substrate is a major factor limiting the growth of heterotrophic bacteria (1). It is likely that bacteria in soil are starved most of the time (2) and only experience infrequent episodes of ample provision of carbon substrate, for example as exudates from a root or organic material released during decay of dead (micro)organisms. Bacteria have developed several strategies to survive extended periods of starvation, such as the development of high-affinity uptake systems to scavenge alternative carbon sources from the surroundings, as well as changes in cell morphology, condensation of the nucleoid and decreased protein synthesis to adapt to a low metabolic activity (3). Many bacteria produce carbon-rich storage materials such as PHA (poly-3-hydroxyalkanoates) and glycogen during periods of substrate availability, which can be utilized to sustain a minimum of metabolic activity when deprived of carbon and energy (4,5).

Several of the microbially mediated processes in the global nitrogen cycle are closely linked to the carbon (C) metabolism of the organisms. One example is denitrification, which is the reduction of NO_3_ ^-^ to N_2_ through anaerobic respiration where the N-oxides are used as terminal electron acceptors when O_2_ becomes scarce. This process can be performed by a diverse range of heterotrophic bacteria, archaea and fungi, which use various forms of organic compounds as electron donors to obtain energy (6) or, in some cases, H_2_ (7). The last step of denitrification is the reduction of N_2_O, a strong climate gas, to harmless N_2_, catalyzed by the N_2_O reductase (Nos) (8,9). It is found in a diverse range of prokaryotic organisms but has not been reported in eukaryotes. Some denitrifying prokaryotes can perform all steps of denitrification, others only some, and lack of the last step is common due to absence of the *nosZ* gene coding for NosZ, or lack of other essential gene(s) in the *nos* operon (10,11), but the amounts of N_2_O released from denitrification in relation to N_2_ (the N_2_O/N_2_ product ratio) is also influenced by transcriptional and post-transcriptional control mechanisms and by various environmental factors (6,12-19).

Denitrification in agricultural soils is a major source of N_2_O, accounting for more than 60% of the global anthropogenic emissions (20,21). A steady increase in atmospheric N_2_O has been recorded since the start of industrialization, largely driven by increasing and excessive use of synthetic fertilizers (22,23) and these emissions are predicted to continue to increase unless novel mitigation options are developed (24,25).

Although the addition of reactive nitrogen compounds via synthetic fertilizers accounts for the main part of the N_2_O emissions from agricultural soil, the N_2_O emitted from legume cropped fields is far from negligible. A compilation of data from ca 70 legume cropped fields estimated ca 1.29 kg N_2_O-N ha^-1^ during one growing season, while the corresponding data for N-fertilized crops and pastures showed emissions of 3.22 kg N_2_O-N ha^-1^ (26). Legumes do not have to rely on uptake of reactive N such as NH_4_ ^+^ or NO_3_ ^-^ but can acquire N through their symbiotic relationship with some groups of bacteria, collectively called rhizobia, that elicit the production of root nodules on the plant in which they fix atmospheric N_2_. In this process, the rhizobia reduce N_2_ to NH_3_ and the plant cells reduce the NH_3_ further to glutamine which they use to produce amino acids and eventually proteins (27). When this N-rich plant material is degraded, organic N is released and mineralized to NH_3_/NH_4_^+^ which will be oxidized to NO_3_ ^-^ by nitrifying organisms. The O_2_ consumption by the nitrifiers, together with the production of NO_3_ ^-^ and the availability of organic compounds from the degraded plants, creates conditions that are conducive to denitrification. A novel approach to minimize N_2_O emissions from agricultural soil is to enhance the populations of N_2_O reducing bacteria (25). In the case of legume crops, which are often inoculated with rhizobia to optimize the N_2_-fixation, there are a few promising studies reporting decreased N_2_O emissions from soybean fields inoculated with rhizobia with the dual capability of efficient N_2_ fixation and N_2_O reduction (28,29).

Based on this, selection of rhizobial strains for development of commercial inoculants should, ideally, take both these aspects into account. One problem is, however, that far from all rhizobia carry the *nosZ* gene that encodes Nos. There are relatively few surveys of denitrification genes in different groups of rhizobia. Complete denitrification, which includes all four reduction steps of NO_3_^-^ to N_2_, has so far mainly been reported for the genus *Bradyrhizobium*, which is the microsymbiont of a range of economically important legume crops such as soybean, cow pea and peanut (30,31). A full set of denitrification reductases in bradyrhizobia include, with few exceptions, the periplasmic NO_3_^-^ reductase Nap; the Cu-containing NO_2_ ^-^ reductase NirK, the *bc*-type NO reductase cNor and a NosZ belonging to clade I (19,32). We recently investigated the denitrification capacity of bradyrhizobia from two strain collections, one obtained from nodules of legume trees and herbs growing in Ethiopia, the other mainly consisting of strains isolated from nodules of peanut growing in China (18,19). In these collections, 50 and 37% of the isolates, respectively, were complete denitrifiers, while the others generally lacked NosZ and thus were potential N_2_O sources. Common to all strains with complete denitrification was a strong preference for N_2_O-over NO_3_^-^ reduction. Transcription analysis and proteomics showed comparable expression levels of Nap and Nos, suggesting a control mechanism at the metabolic level where the electron pathway to Nos, which receives electrons from cytochrome *c* via the *bc1* complex, competes very efficiently with the electron pathway to Nap which goes via the membrane bond NapC.

The results presented in (18,19) were based on experiments with organisms provided with ample amounts of C substrate (electron donor), which is likely to reflect the situation in legume nodules where the microsymbiont receives C from the plant. It can, however, be expected that rhizobia, which may survive for many years in soil (33), spend a large part of their life cycle as free-living organisms in soil where they will experience lack of available C substrate most of the time (2). Rhizobial inoculants that carry NosZ are thus potentially important sinks for N_2_O produced both by themselves and by other soil microbes. It is, however, not known if the competition for electrons favoring N_2_O reduction in well-fed cultures is retained during substrate limitation (“starvation”). Here we exposed cultures of *Bradyrhizobium* strain HAMBI 2125, also studied in (19), to shorter and extended periods of starvation and analyzed the denitrification kinetics, including the electron flow rates to the individual denitrification reductases. We also quantified the cellular abundancies of Nap, Nir and Nos. The results have practical implications, supporting that these organisms can act as sinks for N_2_O under natural conditions. Moreover, the results are ecologically interesting since they show that cultures exposed to extended starvation divided into two opposite denitrification phenotypes, one with very slow metabolism, the other with retained metabolism, possibly reflecting a strategy to increase the chances for survival during periods of starvation.

## Results

### Denitrification kinetics in cultures prepared following Bioassay 1

Two bioassays were developed for starvation experiments, Bioassay 1 and 2 (Figs. 1A and B). In a first experiment, the denitrification gas kinetics, concentrations of NO_3_^-^ and NO_2_^-^ and electron flow rates to the different reductases were compared for well-fed *vs* starved cultures (Figs. 2A-D). For this, the starved cultures were prepared following Bioassay 1 (Fig. 1A), in which the cultures were allowed to synthesize the denitrification reductases in the presence of ample amounts of substrate. Cultures were raised under fully oxic conditions in YMB medium, after which the headspace was made hypoxic to allow transition to anaerobic respiration in response to a gradual depletion of oxygen. After centrifugation/washing, pellets were pooled and inoculated into flasks containing YMB (9.9E+08 cells flask^-1^) or buffer (5.0E+09 cells flask^-1^), supplemented with 1 mM KNO_3_ and 0.25 mM KNO_2_, and with He plus 1 ml N_2_O (around 80 μmol N) in headspace.

**FIG 1.**
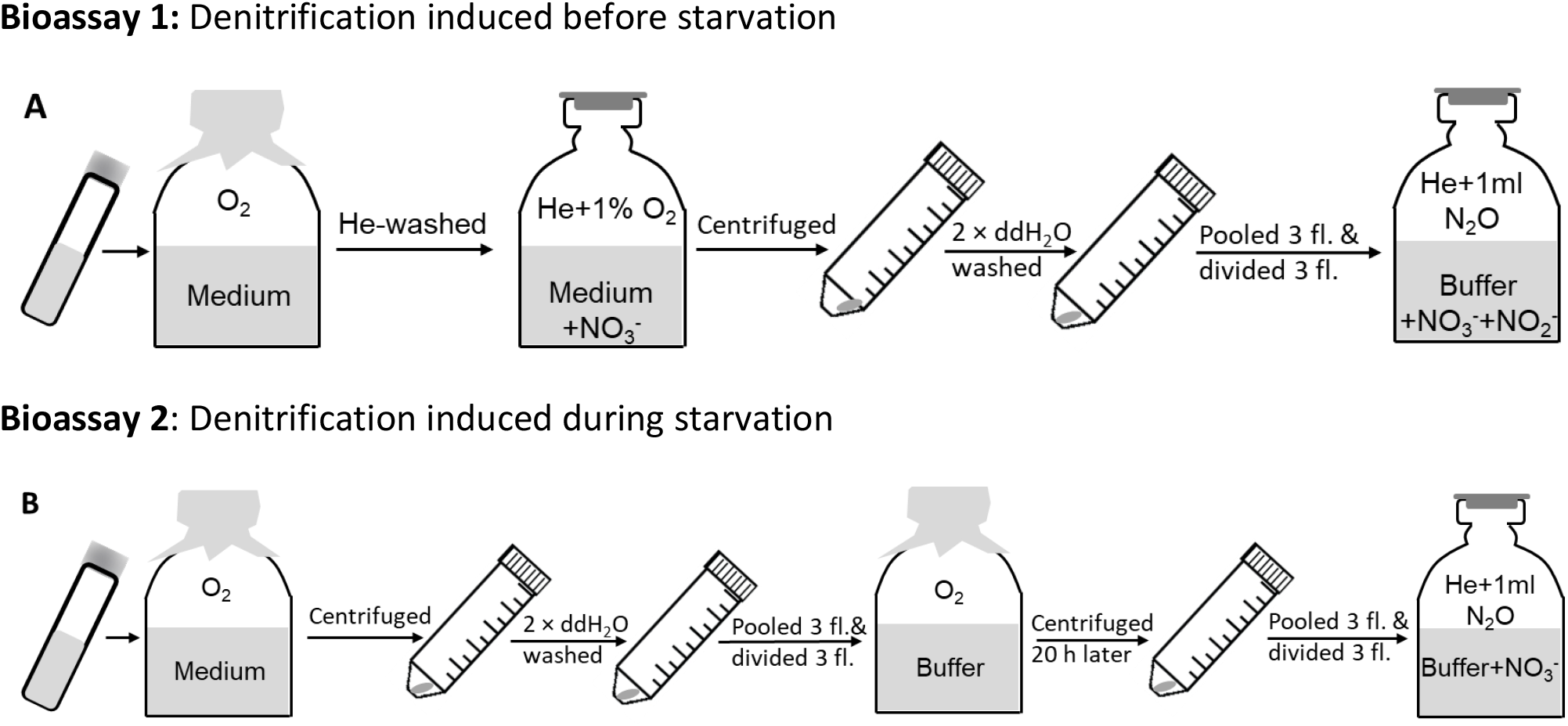
Bioassays for assessing the effect of starvation on electron flow to denitrification enzymes. <u>A: Bioassay 1: starvation of cells with a previously expressed denitrification proteome</u>. Cells were raised from stock cultures in fully oxic flasks containing YMB medium. When OD_600_ reached ∼0.1, the flasks were made anoxic by He-washing, then supplemented with 1% O_2_ in the headspace and 1 mM NO_3_ ^-^ in the liquid. The cultures were allowed to deplete the O_2_ and to initiate denitrification when growing in YMB medium, after which they were centrifuged (10 000 × g at 4 °C for 10 mins) and washed twice in sterile ddH_2_O. The pellets from triplicate flasks were pooled and used to inoculate flasks containing either C-free buffer or YMB medium (well-fed control), in both cases supplemented with 1 mM KNO_3_ and 0.25 or 0.5 mM KNO_2_, and with He and 1 ml N_2_O (around 80 μmol N flask^-1^) in headspace. The starving cells, incubated in buffer, had a low respiratory electron flow rate (mol electrons cell^-1^ h^-1^), initially being 10-18 % that of the well-fed cultures, and then decreasing to reach around 4% after 20 h. Results from experiments using Bioassay 1 are presented in Figs. 2, 3 and 4. <u>B: Bioassay 2: denitrification induced during starvation.</u> Cells were raised from stock cultures in fully oxic flasks containing YMB medium. When OD_600_ reached ∼0.1, the cultures were centrifuged (10 000 × g at 4 °C for 10 mins) and washed twice in sterile ddH_2_O. The pellets from triplicate flasks were pooled, after which they were evenly divided and used to inoculate fully oxic flasks containing C-free buffer. These cultures were incubated for 20 h, then centrifuged after which the pellets were pooled and divided evenly before being inoculated into flasks containing C-free buffer provided with 1 mM KNO_3_, and with He and 1 ml N_2_O (around 80 μmol N flask^-1^) in headspace. The respiration rate (mol electrons cell^-1^ h^-1^) of the starving cultures was 1-4% compared to that of well-fed cultures. All cultures in Bioassays 1 and 2 were incubated at 28°C, and with vigorous stirring (650 rpm). Results from experiments using Bioassay 1 are presented in Figs. 5 and S2.

**FIG 2.**
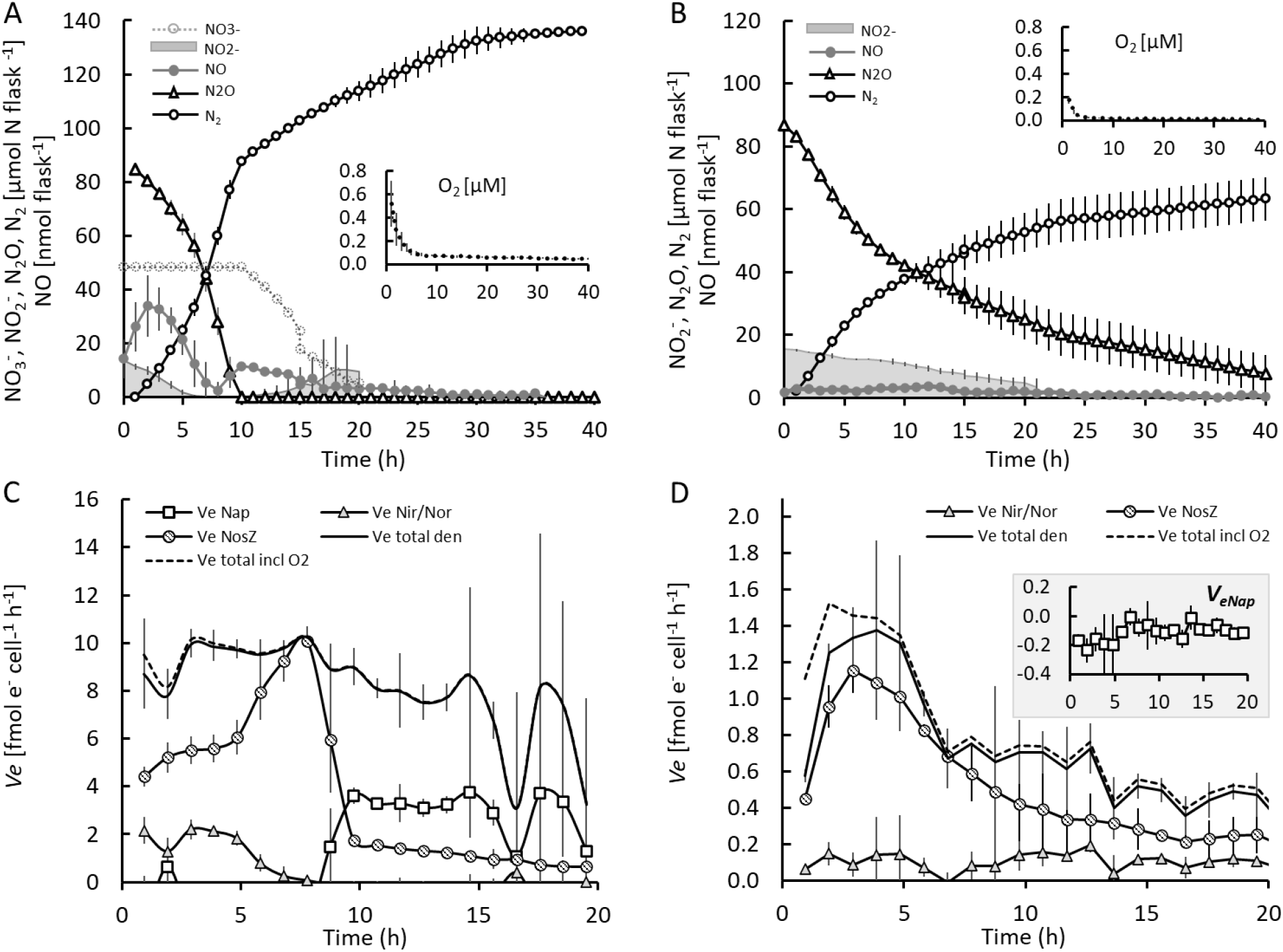
Denitrification kinetics as affected by starvation in cultures with a complete denitrification proteome (Bioassay 1). Cells were allowed to develop a full denitrification proteome under well-fed conditions and were then washed twice in buffer prior to inoculation to flasks with YMB (well-fed control) or buffer (starved), supplemented with 1 mM KNO_3_ and 0.25 mM KNO_2_, and with He plus 1 ml N_2_O (around 80 μmol N) in headspace. A larger inoculum was given to the flasks with buffer (9.9E+08 cells flask^-1^) than to the flasks with YMB (5.0E+09 cells flask^-1^), to secure measurable activity in the starved cells and adequate time resolution of the denitrification kinetics in the well-fed cells. <u>Panels A and B</u> show the denitrification kinetics in well-fed and starved cultures, respectively. The flasks were practically anoxic from the start with an initial O_2_ concentration in the liquid of <0.52 μM which decreased to approximately 0 μM (insets in A & B). <u>Panels C and D</u> show the cell specific electron flow rate. *V*_*e total den*_ designates the total electron flow to the denitrification reductases and *V*_*e total incl O2*_ the total electron flow, including that to denitrification and to O_2_. The electron flow rate to the individual reductases is designated as *V*_*eNap*_, *V*_*eNir*_, *V*_*eNor*_ and *V*_*eNos*_. *V*_*eNir*_ and *V*_*eNor*_ were practically identical and cannot be distinguished from one another in the figure. Inserted panels show *V*_*eNap*_ throughout, including negative values which are due to slight errors in determination of N_2_ and N_2_O (*V*_*eNap*_ was calculated by N-mass balance). Bars in all graphs show standard deviation (n=3).

The initial O_2_ concentration, which was 0.5 μM in cultures with YMB and 0.2 μM in cultures with buffer, was depleted within the first 5 h in both treatments (insets in Figs. 2A and B). The provided NO_2_ ^-^ and N_2_O were reduced simultaneously from the beginning of the incubation in both treatments. The NO_3_ ^-^ was left untouched in the well-fed cultures until the exogenous N_2_O was reduced (Fig. 2A), also seen from lack of electron flow to Nap except for a small peak in electron flow early in the anoxic incubation (Fig. 2C). No NO_3_ ^-^ reduction took place in the starved cultures throughout the entire incubation period, as seen from the lack of electron flow to Nap (Fig. 2D). The negative *V*_*eNap*_ estimated for the starved cultures (inset in Fig. 2D) and the initial phase of the well-fed cultures are probably due to minor errors in calibration of N-gas measurements as well as in parameters used to calculate rates of N-transformation (sampling loss and N_2_-leakage, see Molstad et al. (34)), which amounts to substantial errors in the estimates of NO_3_^-^reduction rates because they are based on N-mass balance (explained in detail by Lim et al. (35)). This implies a relatively high detection limit for NO_3_^-^ reduction, and the negative values cannot be taken as evidence for the complete absence of any electron flow to NO_3_ ^-^. However, the measured NO concentrations lend some support to the claim that *V*_*eNap*_ was ∼0 until depletion of the externally provided N_2_O. In the well-fed culture, the concentration of NO declined to zero in response to depletion of NO_2_ ^-^, and increased soon after, as *V*_*eNap*_ increased (Figs. 2A and C). Likewise, the NO concentration declined to very low values in response to depletion of NO_2_ ^-^ in the starved culture (Fig. 2B).

Fig. 2C and D show the calculated electron flow rates per cell to the individual reductases, and their sum, illustrating their competition for electrons. This shows that the well-fed cells (Fig. 2C) sustained a nearly constant total electron flow rate around 9 fmol e^-^ cell^-1^ h^-1^ throughout, but allocated to different reductases depending on the availability of electron acceptors: As long as NO_2_ ^-^ and N_2_O were both present, Nos captured around 50% of the electrons (*V*_*eNos*_ ∼ *V*_*eNir*_ + *V*_*eNor*_), increasing to 100% when NO_2_^-^ was depleted after 6-7 hours, while the electron flow rate to Nap remained insignificant until the externally provided N_2_O became depleted. The starving cells (Fig. 2D) had an order of magnitude lower total electron flow rate per cell, declining gradually throughout the incubation, and here Nos captured >> 50% of the electrons during the first 10 h (*V*_*eNos*_ >> *V*_*eNir*_ + *V*_*eNor*_), but declined to ∼50% towards the end.

Bioassay 1 was also used in a second experiment, but in this case the cultures were incubated anoxically ([O_2_] <0.21 ±0.04 μM) in buffer containing only one of the N-oxides (NO_3_^-^, NO_2_ ^-^ or N_2_ O) as initial electron acceptor (Figs. 3A-C). This experiment established that starved cultures readily reduced NO_3_ ^-^ from the start when no other N-oxide was provided (Fig. 3A). It also demonstrated that the cell specific electron flow to N-oxides was practically unaffected by the type of electron acceptor provided, except for the slightly lower rates initially for flasks with NO_2_ ^-^. For all treatments, the cell specific electron flow rate decreased gradually throughout, and the levels are very similar to those observed in the first experiment (Fig. 3D).

**FIG 3.**
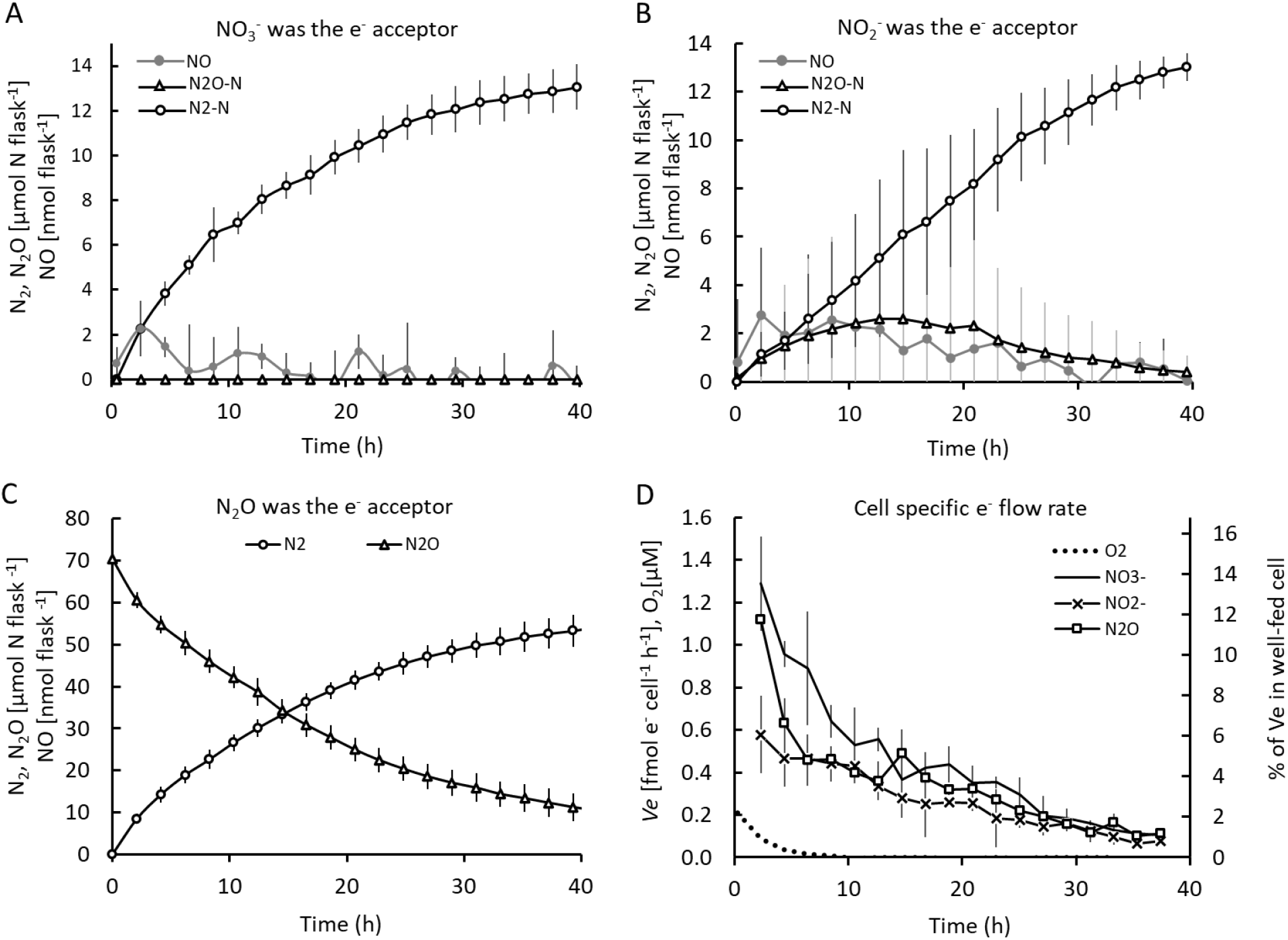
Bioassay 1 with single nitrogen oxides. Experimental conditions as for Fig. 2, but the starving cells (in buffer) were provided with either NO_3_ ^-^, NO_2_ ^-^ or N_2_ O in individual flasks (n=3 for each treatment). <u>Panels A-C</u> show the gas kinetics in flasks provided with 1 mM NO_3_ ^-^ (A), 0.5 mM NO_2_^-^ (B), or 70 μmol N O-N added to the headspace (C). <u>Panel D</u> shows the cell-specific electron flow rate (*V*_*e*_) measured in buffer supplemented with NO_3_^-^, NO_2_^-^ or N_2_O; the O_2_ concentration (left y-axis); and *V*_*e*_ as percentage of the rates in well-fed cultures (right y-axis). The number of cells inoculated into the incubation flasks at the start (0 h of incubation in anoxic buffer) was 3.6E+09 for all treatments. Bars show standard deviation (n=3).

### Cell size and PHA accumulation of starved vs well-fed cultures

The morphologies of single bacterial cells from different treatments (starved/well-fed and anoxic/oxic) were analyzed by phase contrast microscopy (Fig. S1A). By quantifying cell dimensions, we found marginal but statistically significant effects (Mann-Whitney test, p < 0.01) of starvation: while there was no significant difference in cell area of individual cells (Fig. S1B), starved cells were on average ∼12% longer and ∼5% thinner than well-fed cells (Fig. S1B). By further calculating cell volumes, a slight reduction in average cell volume was observed upon starvation of aerobic cells (p=0.057), but not for cells grown under anoxic conditions (Fig. S1C). Furthermore, a qualitative analysis of the presence of PHA granules was done by staining the cells with Nile Red, a lipophilic dye with high affinity for PHA (36). Large foci corresponding to PHA granules were seen in all conditions. It should be noted that, as a lipophilic dye, Nile Red will also bind non-specifically to lipids and membrane, and based on the imaging performed here, it could not be concluded whether there were any significant differences in PHA content between the conditions.

### Denitrification kinetics after providing starving cultures with an artificial electron donor

An experiment was then performed in which TMPD plus ascorbate was added to starved cultures (Bioassay 1), to provide cytochrome *c* with an excess of electrons (37,38). Since cultures treated according to Bioassay 1 were able to provide electrons for denitrification (10-18% of the electron flow in well-fed cells during the first 5 h, then decreasing to ca 4%; Fig. 2D), we expected that the TMPD treatment would reduce or eliminate the oxidation of quinol by the *bc1* complex and that this would allow electrons to flow to Nap via NapC, resulting in NO_3_ ^-^ reduction. We also expected that loading cytochrome *c* with electrons would relieve or weaken the competition for electrons between Nir, Nor and NosZ. The results (Fig. 4) lend little support to the former since the electron flow to Nap remained insignificant until N_2_O had been depleted. But the results confirm an effect of TMPD on the competition between Nos and Nir: *V*_*eNir*_, *V*_*eNor*_ and *V*_*eNos*_ were similar and high (4-5 fmol e^-^ cell^-1^ h^-1^) until NO_2_ ^-^ was depleted. Thus, Nir and Nos competed equally well when the cytochrome *c* pool was fully reduced by TMPD. After depletion of NO_2_ ^-^, *V*_*eNos*_ leaped to its maximum level, 13-14 fmol N cell^-1^ h^-1^, and kept this rate until all the exogenous N_2_O was depleted.

**FIG 4.**
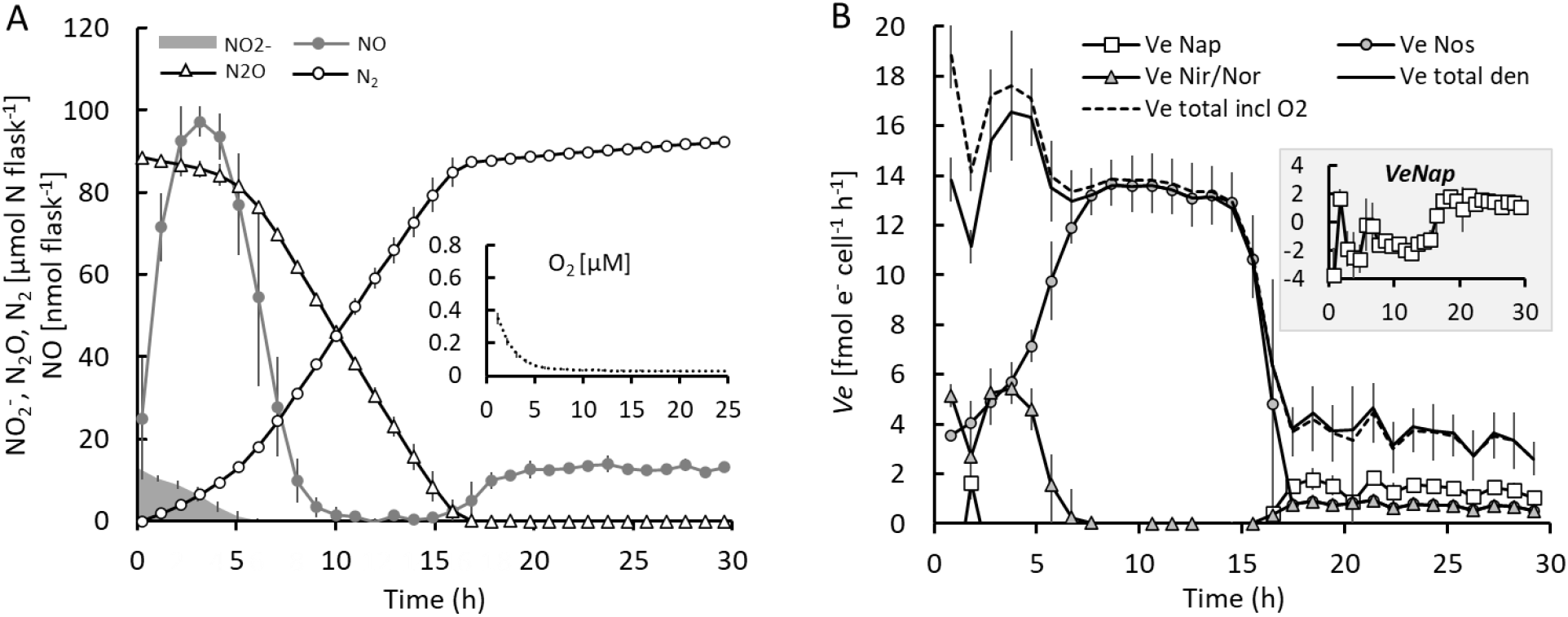
Denitrification in starved cells after addition of TMPD as an external electron donor. Preparation of the cultures followed Bioassay 1 (Fig. 1A), except that 100 μM TMPD and 10 mM ascorbate were injected into the flasks with starving cells 15 min before the first gas sampling. Each flask was inoculated with 5.28E+08 cells (n=4 replicate flasks). The initial O_2_ concentration was 0.35 μM and decreased to approximately 0 within 5 h (inset in A). Bars show standard deviation (n=4). <u>Panel A</u>: kinetics of NO_2_^-^, NO, N_2_O and N_2_ (O_2_ in inserted panel). <u>Panel B:</u> Calculated cell specific electron flow rate (*V*_*e*_, fmol cell^-1^ h^-1^) to each of the reductases Nap, Nir, Nor, and NosZ, and the total electron flow rate (*V*_*etotal*_). The electron flow to Nir was practically identical to the electron flow to Nor (miniscule amounts of NO), and the two are shown as a single graph (*V*_*eNir/Nor*_). Inset plots show *V*_*eNap*_ throughout, including negative values which are due to slight errors in determination of N_2_ and N_2_O (*V*_*eNap*_ was calculated by N-mass balance).

### Denitrification kinetics and reductase abundancies of cultures exposed to extended starvation following Bioassay 2

While Bioassay 1 successfully induced starvation in terms of a downshift in respiration, the rates did not reach a stable level during the assay but declined gradually throughout, apparently approaching more stable low levels after 20 h. On this background, we introduced a more severe starvation assay (Bioassay 2, Fig. 1B) by including a 20 h aerobic incubation in buffer prior to the starvation-denitrification assay, in order to reach a lower and more stable rate of respiration than in the first experiment. In addition, Bioassay 2 was designed to force the cells to synthesize the denitrification proteome while starving. In several preliminary experiments with Bioassay 2 (results not shown), we found a conspicuous variability between flasks regarding the cell specific respiration rate where one or two out of three replicate flasks had four to six times higher cell respiration rates than the other(s). At first, we suspected that it could be due to impurities of flasks or magnets, but meticulous acid washing failed to remove the stochastic variation. Being convinced that the stochasticity reflects a real regulatory switch of the cultures, we performed a final experiment in which 15 replicate flasks were monitored for denitrification kinetics (Fig. 5).

**FIG 5.**
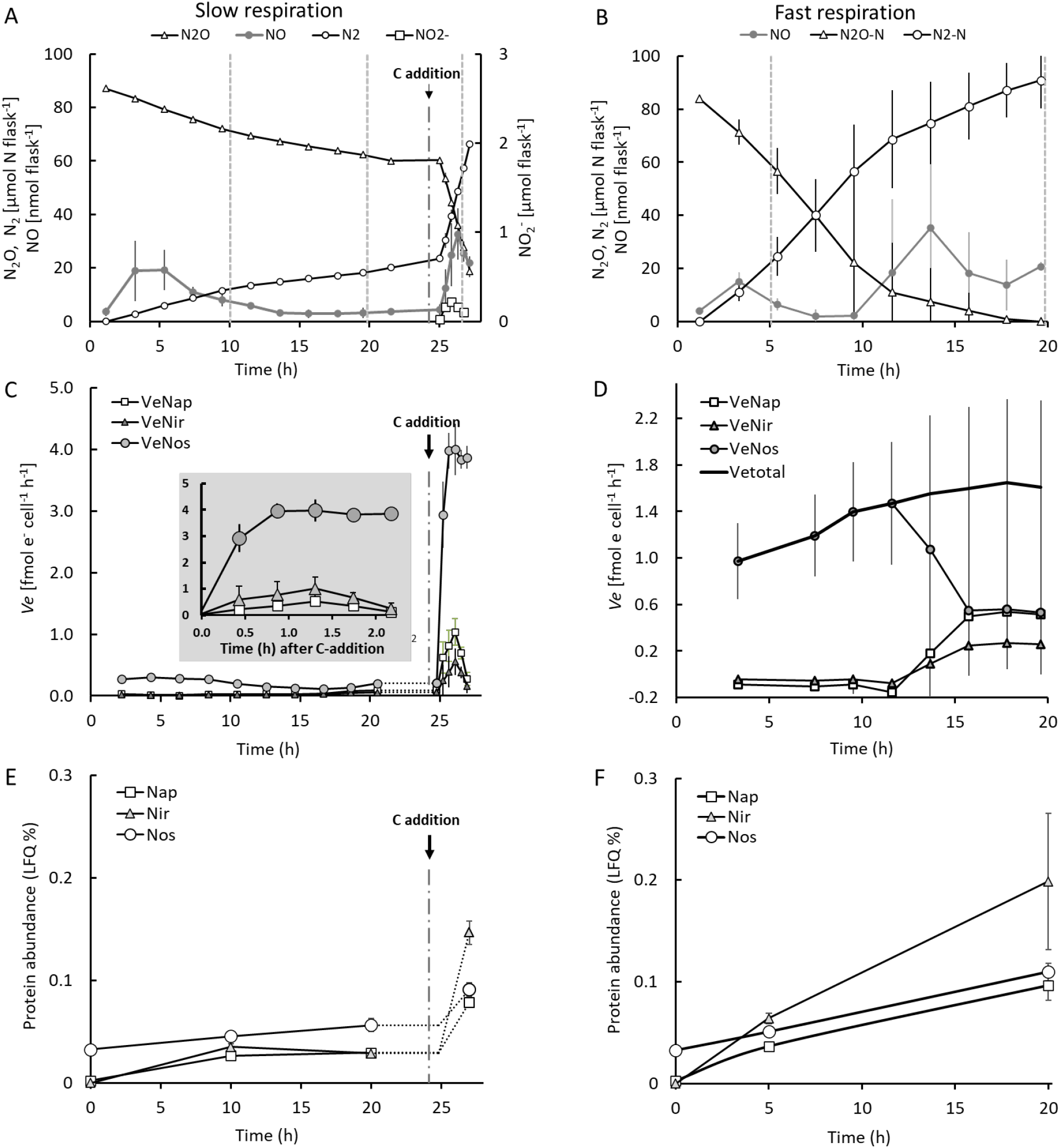
Bioassay 2: stochasticity of starvation response, response to input of organic C and quantification of denitrification enzymes. Altogether fifteen flasks were prepared following Bioassay 2 (see Fig. 1B). All flasks were anoxic (<0.5 μM at the start of the incubation) and contained 50 ml buffer supplemented with 1 mM KNO_3_ and with He plus 1 ml N_2_O in the headspace. The NO_2_^-^ concentrations, measured during the first 5 hours after incubation in the buffer, were approximately 0.59±0.20 μM (not shown). The cultures separated into two distinct phenotypes: nine flasks had “slow” respiration (panel A) and 6 flasks had “fast” respiration (panel B). The electron flow to the individual reductases (fmol e^-^ cell^-1^ h^-1^) for the flasks with slow and fast respiration are shown in panels C and D, respectively. Cultures with “slow” respiration rate had a total electron flow rate of maximum 0.27 e^-^ cell^-1^ h^-1^ (C) and cultures with “fast” respiration rate had a total electron flow rate of 1.0-1.8 fmol e-cell^-1^ h^-1^ (D). The inset plot in panel C shows the electron flow to the individual denitrification reductases after the carbon addition. The cultures (entire flasks, 50 ml) were sampled for proteomics analyses at different time points (panels E and F). At timepoints throughout, marked by dashed vertical lines in panels A and B, three flasks were harvested for proteomics analysis (including 0 h). The six flasks with “fast” respiration were harvested for proteomics analysis at 5 and 20 h of incubation in these anoxic buffers (triplicates at each sampling point). Of the nine flasks with “slow” respiration, three were harvested at 10 and three at 20 h. The remaining three “slow” respiration flasks were supplemented with YMB at 24.8 h, at a concentration which made the buffer a half-strength YMB medium, containing 5 g mannitol l^-1^ and 0.25 yeast extract l^-1^, and then harvested for proteomics at 27 h. The gas measurements and electron flows are averages from the three flasks (n=3) for each phenotype, that were left untouched until the end of the incubation when they were sampled for proteomics. Proteomics analyses were done for triplicate flasks (n=3) at each sampling point. Standard deviations are indicated as bars in all graphs (in several cases not visible due to low variation).

The cultures received initially 1 mM NO ^-^ in the buffer (but no NO_2_ ^-^), and He plus 1 ml N_2_ O (80 μmol N) in headspace. The O_2_ concentration at the time of inoculation was <0.5 μM in the liquid. The flasks separated into two distinct denitrification phenotypes (Figs. 5A and B). Nine flasks showed “low” cell specific respiration rates (total electron flow maximum 0.27 fmol e^-^ cell^-1^ h^-1^), corresponding to approximately 2.7% compared to well-fed cultures (Fig. 5A), while the other six showed a “fast” respiration rate (total electron flow 1.0-1.8 fmol e^-^ cell^-1^ h^-1^). Both phenotypes reduced N_2_O from the beginning of the incubation, showing a strong preference for N_2_O over NO_3_ ^-^. In the flasks with fast respiration, the cells started to reduce NO_3_^-^ in response to depletion of the externally provided N_2_O. In the flasks with slow respiration, N_2_O was not depleted within the time frame of the experiment, and NO_3_ ^-^ reduction remained negligible.

To investigate if cell lysis, and thus release of available C, could explain why some cultures showed the “fast” growth phenotype, we compared the OD_600_ and the number of viable cells in “slow” *vs* “fast” cultures. The samples were taken after incubation for 3.1 h in anoxic buffer, when the two phenotypes were clearly distinct but when some possible growth of cells in the “fast” cultures would not yet hide if lysis had occurred. The OD_600_ spanned from 0.12-0.13 with no statistical difference (p>0.3) between cultures with “fast” and “slow” respiration. Similarly, no difference (p=0.4) was found for the viable counts which showed 1.21E+10 to 1.33E+10 CFU flask^-1^ for the “fast” cultures and 1.23E+10 to 1.34E+10 CFU flask^-1^ for the “slow” cultures (Fig. S2).

We also tested the metabolic integrity of the cells in the flasks with slow respiration by injecting C-substrates (YMB) to the flasks after 24.8 hours, which proved their capacity to quickly regain activity approaching that of well-fed cells (Fig. 5A). To investigate if the observed divergencies in phenotypes were due to differences in denitrification reductase abundancies, we quantified the relative abundances of Nap, NirK and NosZ in samples taken at different time points throughout the incubations (Figs. 5E and F), together with the corresponding denitrification kinetics (Figs. 5A and B) and cell specific electron flow to reductases (Figs. 5C and D). The membrane-bound NO reductase (cNor) could not be extracted quantitatively (the results showed 1000 times lower abundancies than for the other denitrification reductases) and is therefore not shown. The inoculum had been cultured aerobically for 3-4 generations, never permitting the OD_600_ to exceed 0.1, to secure that any denitrification enzymes would be diluted to extinction by aerobic growth, assuming that the transcription of all genes is effectively repressed by oxygen. This strategy was apparently successful since Nap and Nir were undetectable at the start of the anoxic incubation. NosZ, on the other hand, was detected also in the aerobic cultures, suggesting that the *nosZ* gene is constitutively transcribed at low levels in these organisms.

After transferring the cells to anoxic buffer, the abundance of all three reductases increased both in cultures with “slow” and “fast” respiration. Cultures with “slow” respiration rate synthesized less denitrification reductases than those with “fast” respiration rate during the first 20 h. The relative abundances of the different reductases also differed between the two phenotypes. In cultures with “slow” respiration, NosZ was significantly more abundant than Nap and Nir (p<0.01), which were comparable (Fig. 5E). Cultures with “fast” respiration instead contained higher abundancies of NosZ and Nir compared to Nap at 5 h. After this, the abundance of Nir increased more than that of the others and at 20 h the LFQ of Nir was 0.20 ± 0.07, while the abundancies of NosZ and Nap were approximately half of that (Fig. 5F). After addition of carbon substrate to the cultures with “slow” respiration, a rapid synthesis of all three reductases took place. This synthesis was most prominent for Nir which, after a couple of hours, had increased three-fold, reaching a relative abundance that was twice as high as Nap and NosZ (Fig. 5E), resembling the abundance profile of the cultures with “fast” respiration (Fig. 5F).

## Discussion

Detailed eco-physiological studies during the past decades have revealed several aspects of how the denitrification process is regulated in different organisms (6,13,39). Most of this knowledge is, however, based on laboratory studies where cultures have been grown under optimal conditions, while less attention has been paid to understanding how denitrification, and particularly the release of N_2_O, is affected if cells are starved, which is the normal state of cells in most natural environments. We focused on how starvation, i. e. lack of carbon substrate, affects denitrification and N_2_O release. The general notion has been that N_2_O reductase is less competitive for electrons than the other denitrification reductases, leading to emissions of N_2_O when the availability of electron donors is low. This is largely based on a single study of *Alcaligenes faecalis* (40) and has been supported by some studies of complex communities but contested by others (41-43). *A. faecalis* can perform partial denitrification reducing NO_2_ ^-^ to N_2_ using NirK, cNor and NosZ clade I, but this organism lacks dissimilatory NO_3_ ^-^ reductases (Nar or Nap). The study by Schalk-Otto (40), performed in continuous cultures, showed increased N_2_O release under low substrate concentrations, and it was suggested that N_2_O reductase did not compete successfully with the other reductases for electrons from cytochrome *c*, possibly due to lower affinity for the electron donor. This conclusion needs, however, further verification by more detailed studies of the mechanism involved. Moreover, such studies need to be extended to other groups of denitrifying microorganisms and should include organisms carrying a complete denitrification pathway (thus with Nar and/or Nap). Research over the past decades has revealed diverse denitrification phenotypes among even closely related bacteria, with implications for their accumulation of denitrification intermediate products (15,44).

The phenotype described for a range of taxonomically diverse *Bradyrhizobium* strains with a complete denitrification pathway is characterized by a strong preference for N_2_O over NO_3_^-^ when grown in full-strength YMB medium under denitrifying conditions (18,19). The present study provides compelling evidence that this preference is retained when the organisms are starved for carbon and energy, with practically no electron flow to Nap when N_2_O was present (Figs. 2A-D). This was not due to too low levels of Nap since the cultures showed well-functioning Nap activity if NO_3_^-^ was provided as the only initial electron acceptor (Fig. 3A). Further evidence for this was obtained from the proteomics results, which showed that the cultures, even when grown under more severe starvation conditions using Bioassay 2, were able to produce comparable amounts of Nap, Nir and Nos (Figs. 5E and F). Therefore, the lack of NO_3_^-^ reduction during the period when there was N_2_O in the system could not be explained by a lack of Nap molecules. Instead, the results suggest the same metabolic-level phenomenon as found for well-fed cultures in this study (Fig. 2A) and earlier (18,19), where NosZ outcompetes Nap for electrons, leaving Nap virtually without electrons so long as exogenous N_2_O is available.

The attempt to tweak the electron flow toward Nap by the addition of TMPD and ascorbate did not result in measurable NO_3_^-^ reduction in the starved cultures (Fig. 4A). A recent study by Mania et al. (18) demonstrated that well-fed *Bradyrhizobium* cells did reduce some NO_3_ ^−^ in the presence of N_2_O, if provided with TMPD and ascorbate. This suggested that NosZ (as well as Nir and Nor) received electrons from strongly reduced cytochrome *c*, thus bypassing the electron flow via the *bc1* complex, which allowed Nap to receive electrons from quinol, delivered from the TCA cycle. The specificity of the electron delivery from TMPD to cytochrome *c* cannot be taken for granted, however, and the result by Mania et al. (18) could instead reflect a minimum of electron flow from TPMD to quinon or to NapC, directly, or indirectly. In the present experiment with starved cells, TMPD + ascorbate failed to induce measurable electron flow to Nap in the presence of N_2_O (Fig. 4B). This probably reflects the marginal electron flow from the TCA cycle to the quinon/quinol pool due to starvation. A separate experiment supported this, showing that when carbon substrate (YMB) was added to starved cultures, this provided enough electrons to support some Nap activity, although NosZ activity still dominated (Fig. S3).

At the same time, the result refutes the concerns regarding unspecific electron delivery of electrons from TMPD to quinon. Functional Nap was apparently present, and NO_3_^-^ reduction started when N_2_O was almost depleted, with *V*_*eNap*_ being 2 fmol e^-^ cell^-1^ h^-1^ which was twice as high as *V*_*eNir/Nor*_ (Fig. 4B). The latter is as expected, since the reduction of 1 mole of NO_3_^-^ to NO_2_^-^ requires 2 mole electrons, while reduction of NO_2_^-^ and NO requires 1 mole electrons. Furthermore, the electron flow to NosZ was the same as to Nir and Nor, suggesting that Nir and NosZ competed equally well for electrons from cytochrome *c*. The maximum total electron flow of 13-14 fmol e^-^ cell^-1^ h^-1^ when the cytochrome *c* pool was saturated with electrons from TMPD is likely to be close to the maximum capacity of the denitrification pathway. This electron flow rate is higher than the total electron flow rate in the well-fed cells, which was 8-10 fmol e^-^ cell^-1^ h^-1^ (Fig. 2C). The factor limiting *V*_*eNos*_ to 13-14 fmol cell^-1^ h^-1^ is plausibly the rate of electron delivery from cytochrome *c* and/or *k*_*cat*_ for Nos.

In another experiment we added YMB medium to “slow respiration” phenotypes of the cultures that had been exposed to extended starvation (Figs. 5A, C and E). These cultures had some NosZ activity (*V*_*eNos*_ 0.1-0.3 fmol e^-^ cell^-1^ h^-1^), while Nap and Nir activities could not be detected. Addition of the electron donor (YMB) led to an immediate upshoot in the activities of all reductases. This could not have been the case if the reductases were not present already, which was proven by the proteomics results. Taken together, the results support that the absolute preference for N_2_O over NO_3_ ^-^ was due to competition for electrons, also under severe starvation conditions.

It is well known that bacteria have developed a range of physiological responses to tackle starvation. Some survive by forming spores, but most bacteria survive by strongly reduced metabolic rates and minimizing the synthesis of some proteins while upregulating others such as genes for high-affinity transporters, essential repair mechanisms and alternative energy sources (3,45,46). Some such changes have been observed in rhizobia belonging to *Rhizobium leguminosarum*, which stayed viable for long periods (55 days) of C starvation (47). Changes in cell size are common during long-term starvation, sometimes leading to the formation of small or even ultra-microcells which may have increased tolerance to antibiotics and other stresses (2,46). Another way for many bacteria, including rhizobia, to survive is to use carbon stored as polyhydroxyalkanoates (PHA) or glycogen, formed during periods of ample nutrient abundance (48). In the present study, the metabolic activity, measured as anaerobic respiration rate, decreased to between 1 and 18% that of well-fed cultures, depending on which starvation bioassay was used. We did not, however, detect any obvious decrease in cell size when comparing cells starved for 24 h, although cell morphologies were slightly altered (Figs. S1A and B). PHA was observed in the starved cells as well as well-fed ones, as shown with Nile red staining (Fig. S1A) and no obvious reduction in PHA could be observed during starvation from our assays. Since this was a comparably short period of starvation, it is conceivable that the cells would make use of the stored carbon if the starvation was prolonged. It could be speculated that one reason for not using the PHA reserves (and glycogen) immediately, is that these are saved to be used during bacteroid formation (48). To clarify these issues, more detailed studies are however needed.

A striking separation into two phenotypes during starvation, as shown in Fig. 5, was observed in repeated experiments (Bioassay 2), each time with about two thirds of the flasks showing a “fast” and the others “slow” respiration: 1.0-1.8 fmol e^-^ cell^-1^ h^-1^ vs 0.1-0.3 fmol e^-^ cell^-1^ h^-1^, respectively. Phenotypic heterogeneity has been observed in single-strain cultures of various bacteria when exposed to carbon substrate deficiency and may, or may not, be due to mutations during starvation for one week or more (3,49). Mutations in the entire population in several replicate flasks are unlikely and cannot, however, have caused the rapid diversification into “slow”- or “fast” respiration in the present study. It may instead reflect a stochastic phenomenon, or that the culture contained different subpopulations. We speculated that the “fast respiration” phenotype may be due to a fraction of the cells dying, allowing the other cells to survive on nutrients released from lysed cells, as seen for other bacteria (3). However, this would require lysis of a substantial fraction of the cells, which is refuted by the observation that the flasks with “slow” and “fast” phenotypes had practically identical numbers of viable cells (Fig. S2). Thus, further studies are needed to understand this phenomenon of different respiration rates.

Although the starvation bioassays developed for this study cannot be regarded as a close mimicking of the conditions in natural environments, it is conceivable from the experiments that these organisms are potentially strong sinks for N_2_O when living in soil under fluctuating availability of carbon substrate. Bioassay 1 is probably closer to a “real-world” situation than Bioassay 2, since it is likely that denitrifying bacteria experience regular fluctuations in oxygen and thus are not devoid of denitrification reductases if they enter starvation. On the other hand, Bioassay 2, where the cells had to produce the denitrification proteome in the absence of external electron donors (C-substrate), showed that even under these conditions N_2_O reduction strongly dominated over NO_3_^-^ reduction.

## Materials and methods

### Bacterial strain and culture preparations

*Bradyrhizobium* strain HAMBI 2125, originally isolated from nodules of *Arachis hypogaea* growing in Sichuan, China (50), was used in all experiments. This strain, which is closely related to *Bradyrhizobium ottawaense*, contains the genetic set-up for complete denitrification (19). A culture was raised from one single colony after streaking on agar plates. After checking the purity by sequencing the 16S rRNA gene (19), portions were preserved in 15% glycerol at -80 °C. Cultures for all the experiments were started from the -80 °C stocks and raised under fully oxic conditions in 120 ml serum flasks containing 50 ml Yeast Mannitol Broth (YMB): 10 g l^-1^ D-Mannitol, 0.5 g l^-1^ K_2_HPO_4_, 0.2 g l^-1^ MgSO_4_·7H_2_O, 0.1 g l^-1^ NaCl and 0.5 g l^-1^ yeast extract (51). All incubations were done at 28 °C. Medical flasks (120 ml) were used in all experiments. A magnet in each flask secured vigorous stirring (600-700 rpm) to avoid cell aggregation and to optimize the gas exchange between liquid and gas phases (18). To prevent that the cells experienced anoxia during this oxic incubation, and thus to avoid de novo synthesis of denitrification reductases, portions were regularly transferred to new flasks containing fresh medium, so that the OD_600_ was never allowed to exceed 0.1 (19). These aerobically grown cultures were used as inoculants in the “starvation bioassays” described below.

Flasks for cultures incubated under hypoxic conditions were prepared as described in (18). Briefly, flasks (120 ml) containing 50 ml buffer (or medium) were capped with sterilized, gas tight butyl rubber septa (Matrix AS, Norway). The air was removed by applying vacuum repeatedly (6 × 360 s) and He was then filled for 30 s, after which the overpressure was released. The flasks were left for two days to equilibrate the gases between the headspace and liquid (52). Then, 0.7 ml or 1 ml O_2_ (equal to 1 or 1.5 vol %) and 1 ml N_2_O (70 to ∼80 μmol N flask^-1^) was injected into the headspace and sterile filtered solutions of KNO_3_ (and sometimes KNO_2_) were added to the liquid reaching initial, desired concentrations (0.25 -1 mM).

### Starvation bioassays

Starvation bioassays were established, which followed one of the procedures described in Fig. 1. In Bioassay 1 “Mild starvation”, cultures were allowed to make the transition to denitrification in full-strength YMB before being exposed to starvation. Oxically grown, well-fed cultures were incubated for 48 h after which the headspace was replaced by He and 1% O_2_, and 1 mM KNO_3_ was added to the medium. When O_2_ was depleted, the cultures were centrifuged (10 000 × g at 4 °C for 10 min) and washed twice in sterile ddH_2_O. The pellets (triplicates) were pooled to reduce bias in the form of variations due to the centrifugation/washing. Each pellet was divided into three and used to inoculate anoxic flasks containing 50 ml C-free buffer supplemented with 1 mM KNO_3_ and 0.25 or 0.5 mM KNO_2_ and with He and/or 1 ml N_2_O (around 80 μmol N flask^-1^) in the headspace. When incubated in the buffer, the respiration rate of the cultures was 10-18% that of well-fed cultures during the first 15 h, then it decreased to about 4%.

In Bioassay 2 (“extended starvation”), denitrification was instead induced after having exposed the cells to starvation for 20 h. The cultures were raised in fully oxic flasks containing YMB medium. When OD_600_ reached ∼0.1, the cultures were centrifuged (10 000 × g at 4 °C for 10 mins) and washed twice in sterile ddH_2_O. The pellets (triplicates) were pooled after which they were evenly divided and used to inoculate fully oxic flask containing C-free “starvation buffer”. These cultures were incubated for 20 h, then centrifuged after which the pellets were pooled and divided evenly before being inoculated into flasks containing C-free “starvation buffer” with 1 mM KNO_3_, and with He and 1 ml N_2_O (around 80 μmol N flask^-1^) in headspace. The respiration rate of cultures exposed to “extended starvation” was ca 1-4% compared to that of well-fed cultures. Results from the different starvation bioassays were compared to those from well-fed cultures. To avoid biases, the treatments, which were in all cases set up at least in triplicates (n ≥ 3), were the same regarding centrifugations and washings until the last step when pooled cells were inoculated to anoxic flasks containing either YMB or buffer. The carbon source in the YMB medium comprised >200 times surplus of electron donor compared to electron acceptors throughout all incubations, thus ensuring that the electron donor was not depleted. All cultures were incubated at 28°C, and with vigorous stirring (600-700 rpm).

### Addition of YMB medium or TMPD as electron donors to starved cultures

Experiments were performed to investigate how starved cultures of *Bradyrhizobium* strain HAMBI 2125 responded to the addition of an electron donor, either provided as an artificial electron donor (Fig. 4) or as YMB medium (2 ml mannitol /yeast solution providing them with a substrate concentration corresponding to half-strength YMB, thus 5 g mannitol and 0.25 g yeast l^-1^, Fig. 5). As artificial electron donor we used sodium TMPD (N, N, N’, N’-tetramethyl-p-phenylenediamine) which, in the presence of ascorbate, donates electrons to cytochrome c thus providing electrons to Nir and NosZ (18,37). Ascorbate and TMPD (both from Sigma-Aldrich®, Germany) were dissolved in ddH_2_O or 96% ethanol, respectively, and filter sterilized. The solutions were added to the incubation flasks 10-15 mins before gas sampling. The effect of different concentrations of TMPD (100, 250, and 500 μM in the culture buffer) combined with 10 mM ascorbate on N_2_O reduction was checked prior to the main experiments. This showed that 500 μM TMPD had an obvious inhibition effect on N_2_O reduction, while 100 μM and 250 μM showed no inhibitory effect (not shown). To minimize other, possible effects we used 100 μM TMPD for the experiments (as done in Mania et al. (18)). In another experiment, shown in Fig. S3, 4 ml of a mannitol/yeast solution was added to flasks containing 50 ml starved cultures, providing them with a substrate concentration corresponding to full strength YMB (10 g mannitol and 0.5 g yeast l^-1^).

### Monitoring of gas kinetics, NO_3_ ^-^ and NO_2_ ^-^ concentrations and electron flow rates

The culture flasks were placed in a robotized incubation system and the headspace gas was sampled frequently for N_2_, N_2_O, NO and O_2_ measurements (34). Gas losses caused by sampling were taken into account when calculating the production and consumption of gases, as described by Molstad et al. (34) and Mania et al. (18). The concentration of O_2_ in the anoxic incubation flasks was <0.6 μM at the start of the experiment, which was well below the level for initiating denitrification (4.6 μM; (19)).

The NO_2_ ^-^ concentrations were monitored as described in Mania et al. (18). Briefly, samples (0.1-0.5 ml) were taken every 1 or 2 hours (n=3) from the liquid phase through the septum of the flasks using a sterile syringe. To avoid that the gas kinetics was affected by the sampling, a set of flasks was dedicated to NO_2_ ^-^ measurements and a parallel set was left untouched for gas measurements. NO_2_ ^-^ was determined using a chemoluminescence NOx analyzer (Sievers NOA™ 280i, GE Analytical Instruments) after at first reducing the NO_2_ ^-^ to NO by adding 10 μl liquid sample into a purging device containing a reducing agent (50% acetic acid with 1% (w/v) NaI) (53,54). The NO_2_ ^-^ concentrations were determined against a standard curve (range 0-2 mM NO_2_ ^-^; r^2^= 0.999).

The cell specific electron flow rates (*V*_*e*_, mol e^-^ cell^-1^ h^-1^) for each time increment between two gas samplings were calculated as *V*_*eflask*_(t)/(*N(0)+E*_*cum*_*(t)*Y*), where *V*_*eflask*_*(t)* is the electron flow rate in the flask (mol e^-^ flask^-1^ h^-1^) calculated from measurements, *N(0)* is number of cells in the flask at time=0, *E*_*cum*_*(t)* is the cumulated electron flow (mol e^-^ flask^-1^) at time=t, and Y is the yield per mol electron (cells mol^-1^ e^-^) for HAMBI 2125 as measured previously (19). The conversion factor 5.8E8 cells ml^−1^ *OD_600_ ^−1^ was used to convert OD_600_ to cell numbers (19).

### Proteomics

The abundances of Nap, Nir and NosZ were quantified in starved cultures treated as described for Bioassay 2 (extended starvation). Altogether eighteen flasks were prepared according to Fig. 1B. After the 20 h aerobic incubation in C-free buffer, three flasks were harvested for proteomics analysis and, following centrifugation/washing, the three cell pellets from these flasks were frozen individually at -20 °C. The cultures in the other flasks were pooled three by three and after centrifugation/washing, each cell pellet was divided in three and used to inoculate new flasks containing C-free buffer with NO_3_ ^-^ in the liquid and with He and 1 ml N_2_O in the headspace (Fig. 1B). These flasks (fifteen in total) were placed in the robotic incubation system for monitoring of gas kinetics and NO_2_ ^-^ concentrations as described above. The entire culture volume (50 ml) was harvested from each of triplicate flasks at different time points during the anoxic incubation. The cultures in six of the flasks showed a “fast” respiration rate (total electron flow 1.0-1.8 e^-^ cell^-1^ h^-1^) and were harvested at 5 and 20 h of incubation in anoxic buffer (triplicates at each sampling point). The cultures in the other nine flasks showed a “slow” respiration rate (maximum total electron flow rate was 0.27 fmol e^-^ cell^-1^ h^-1^). Six of them were harvested in triplicates after 10 and 20 h in anoxic buffer. The remaining three flasks from the cultures with “slow” respiration rate received a portion of YMB at 24.8 hand were harvested at 27 h. Harvested cell cultures were centrifuged and stored as pellets at -20 °C. The protein extraction was as described in Gao et al. (19). Briefly, the thawed cell pellets were resuspended in lysis buffer (20 mM Tris–HCl pH 8, 0.1% v/v Triton X-100, 200 mM NaCl, 1 mM DTT). They were then subjected to bead beating (3 × 45 s) with glass beads (particle size ≤ 106 μm; Sigma) using a MP Biomedicals™ FastPrep-24™ (Thermo Fischer Scientific) at maximum power and with cooling on ice between the cycles. After centrifugation to remove cell debris (10 000 × g; 5 min), the supernatant, containing water soluble proteins, was used for proteomic analysis using an Orbitrap mass spectrometer (described in (19,55)). Quantification was based on LFQ (label-free quantification) in MaxQuant (56), and the relative abundance of the individual reductases was calculated as percentages of the sum of all protein abundances for each time point. The nitric oxide reductases NorB/C were not measured since only a small fraction of these membrane-bound enzymes can be accurately obtained with this extraction protocol (approximately 1/1000^th^ the quantity of the other reductases).

### Viable counts and microscopy

The number of viable cells in starved cultures showing slow and fast respiration rates, observed in flasks prepared according to Bioassay 2, was determined by plating dilutions of the cultures on yeast-mannitol agar (YMA). The morphology of cells from well-fed cultures and starved cultures was compared using phase contrast microscopy and a qualitative determination (presence/absence) of PHA was done by staining with Nile Red (Sigma-Aldrich) followed by fluorescence microscopy. Microscopy was performed on a Zeiss AxioObserver with an Orca-Flash4.0 CMOS camera (Hamamatsu Photonics) controlled by the ZEN Blue software. Images were taken with a 100x phase contrast objective. A HPX-120 Illuminator was used as light source for fluorescence microscopy. Images were prepared using ImageJ and analysis of cell sizes was done using the ImageJ-plugin MicrobeJ (57).

## Acknowledgment

This project was supported by Kingenta Ecological Engineering Group Co., LTD and by the project PASUSI financed by the EC Horizon2020 ERA-NET Cofund Programme, Grant agreement No. 727715 and by the Research Council of Norway, projects No. 290488 and 325770. Yuan Gao is grateful to the China Scholarship Council (CSC) for financial support. We thank Gro Stamsås for help with microscopy.

