## Supplemental Figure S1 for "Denitrification by bradyrhizobia under feast and famine and the role of the bc1 complex in securing electrons for N_2_O reduction"

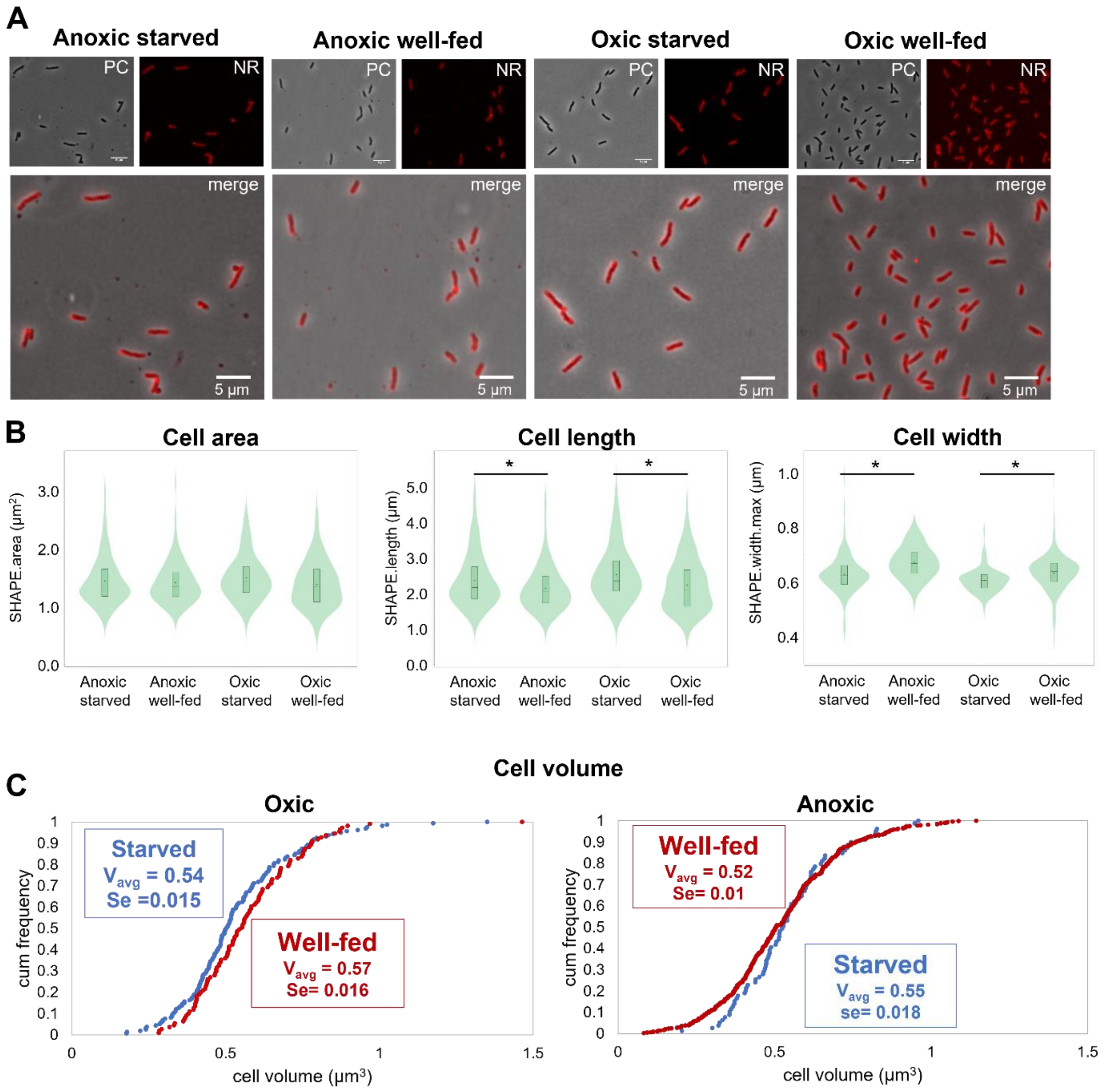


**FIG S1** Cell size, morphology and PHA content of *Bradyrhizobium* strain in carbon optimal or carbon starvation condition. To investigate the effect of carbon source and $O_2$ on the cell morphology, *Bradyrhizobium* strain HAMBI 2125 was raised from stocks in flasks containing YMB medium ("well-fed") or buffer ("starved") with $O_2$ or N-oxide ($NO_3^-$ or $NO_3^-$ plus $N_2O$) as electron acceptors. The cultures were incubated at 28 °C with vigorous stirring (600 rpm using a magnetic stirrer) for five days. To secure that the $OD_{600}$ did not exceed 0.1, portions of the culture were regularly inoculated into new flasks. After 5 days, 1 ml volumes of the cultures were taken and fixed in 1 ml of a 1:1 mixture of paraformaldehyde (2% wt/vol) and glutaraldehyde (2.5% vol/vol). The samples were fixed for 1 hour at room temperature and stored over night at 4°C, then observed under microscopy directly or after staining with Nile red. Samples from **oxic well-fed cultures** were taken as control. Other treatments were prepared as follows: **Anoxic well-fed cultures** were prepared following Bioassay 1 (see Fig. 1A, main text). Briefly, flasks containing oxic, well-fed cultures (in YMB medium) with $OD_{600}$ <0.1 were covered by septa and replacing the headspace air with He, 1 mM $NO_3^-$ and 0.7 ml $O_2$ were added as electron acceptors. Samples were taken after three days of incubation, when the cultures were transitioning from aerobic to anaerobic respiration and thus contained the whole set of denitrifying enzymes. **Oxic starved cultures** were prepared following Bioassay 2 (Fig. 1B in main text), i. e. cultures incubated oxically in YMB were centrifuged and the cell pellets were washed twice using autoclaved $ddH_2O$, then added into oxic flasks containing C-free buffer. Samples were taken after 20 h incubation. **Anoxic starved cultures** were prepared following Bioassay 2, i. e. oxically incubated, starved cultures were centrifuged and added into pre-prepared anoxic flasks containing C-free buffer with 1 mM $NO_3^-$ and with 1% $N_2O$ in the headspace. These cultures, which had to synthesize the denitrifying enzymes during carbon starvation, were sampled for microscopic analysis after 3 h of incubation under anoxic, starved conditions. Panel A: Phase contrast (PC), fluorescence (Nile Red, NR) and the merged images of representative cells are shown for all four conditions. PHA is shown as large Nile Red foci within the cells. Panel B: Cell area, cell lengths and cell widths are displayed as violin plots. More than 75 cells were measured for each condition. Asterisks indicate significant differences between samples (Mann-Whitney test, $p < 0.01$). Starvation had a statistically significant effect on both length (increasing) and width (decreasing). Panel C: Distribution of cell volumes (as calculated from width and length of individual cells), and average cell volumes for each treatment as inserts in the panels. Statistically, the effect of starvation on the average cell volume was barely significant for aerobic cells ($p=0.057$), and not for cells grown under anoxic conditions ($p=0.13$).
