## Supplemental Figure S2 for "Denitrification by bradyrhizobia under feast and famine and the role of the bc1 complex in securing electrons for N_2_O reduction"

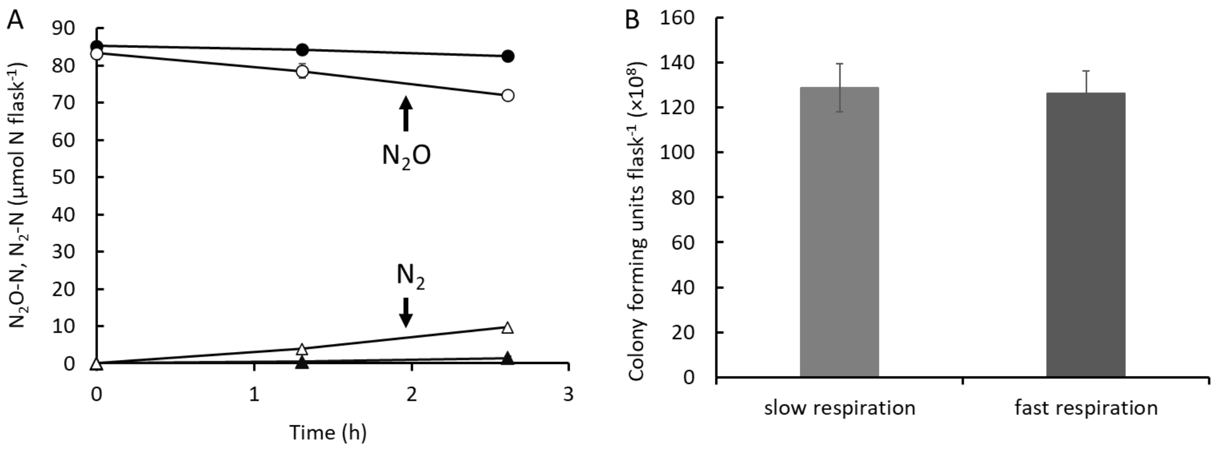


**Fig S2** Nitrous oxide reduction and viable counts of cultures with slow *vs* fast respiration rate after exposure to extended starvation (Bioassay 2). Twelve cultures were prepared following Bioasssay 2 (see Fig. 1B in the main text). Triplicate cell pellets were pooled and then divided into three new flasks containing carbon-free buffer with 1 mM $NO_3^-$and with 1ml $N_2O$ in the headspace. The cultures in four of the flasks showed a "fast" respiration rate ($N_2$ production rate was 2.51 ±0.86 µmol N $flask^{-1}$; n=4), while the remaining cultures had a "slow" respiration rate ($N_2$ production rate was 0.43 ±0.10 µmol N $flask^{-1}$; n=8). Three flasks of each phenotype were chosen for viable counts to determine if cell lysis may have occurred in the cultures with "fast" respiration. **A.** Gas measurements for three flasks of each phenotype ("slow" or "fast" respiration) from which samples for viable counts were taken. $N_2O$ reduction (circles) and $N_2$ production (triangles) in "fast" (open symbols) and "slow" cultures (filled symbols). Bars represent standard deviation (n=3) but are in most cases too small to be visible. **B.** Viable counts (colony forming units; CFU). Samples (1 ml) were taken from each flask after 3.1 h of anoxic incubation in C-free buffer (last step of the assay). The $OD_{600}$ at this time point was not significantly different between the cultures with fast vs slow respiration rate ($P > 0.3$). Diluted samples were streaked on YMA (yeast mannitol agar) plates. No difference in CFU $flask^{-1}$ was found between the cultures with fast respiration rate and slow respiration rate (P=0.4). The bars represent standard deviation (n=3).
