## Supplemental Figure S3 for "Denitrification by bradyrhizobia under feast and famine and the role of the bc1 complex in securing electrons for N_2_O reduction"

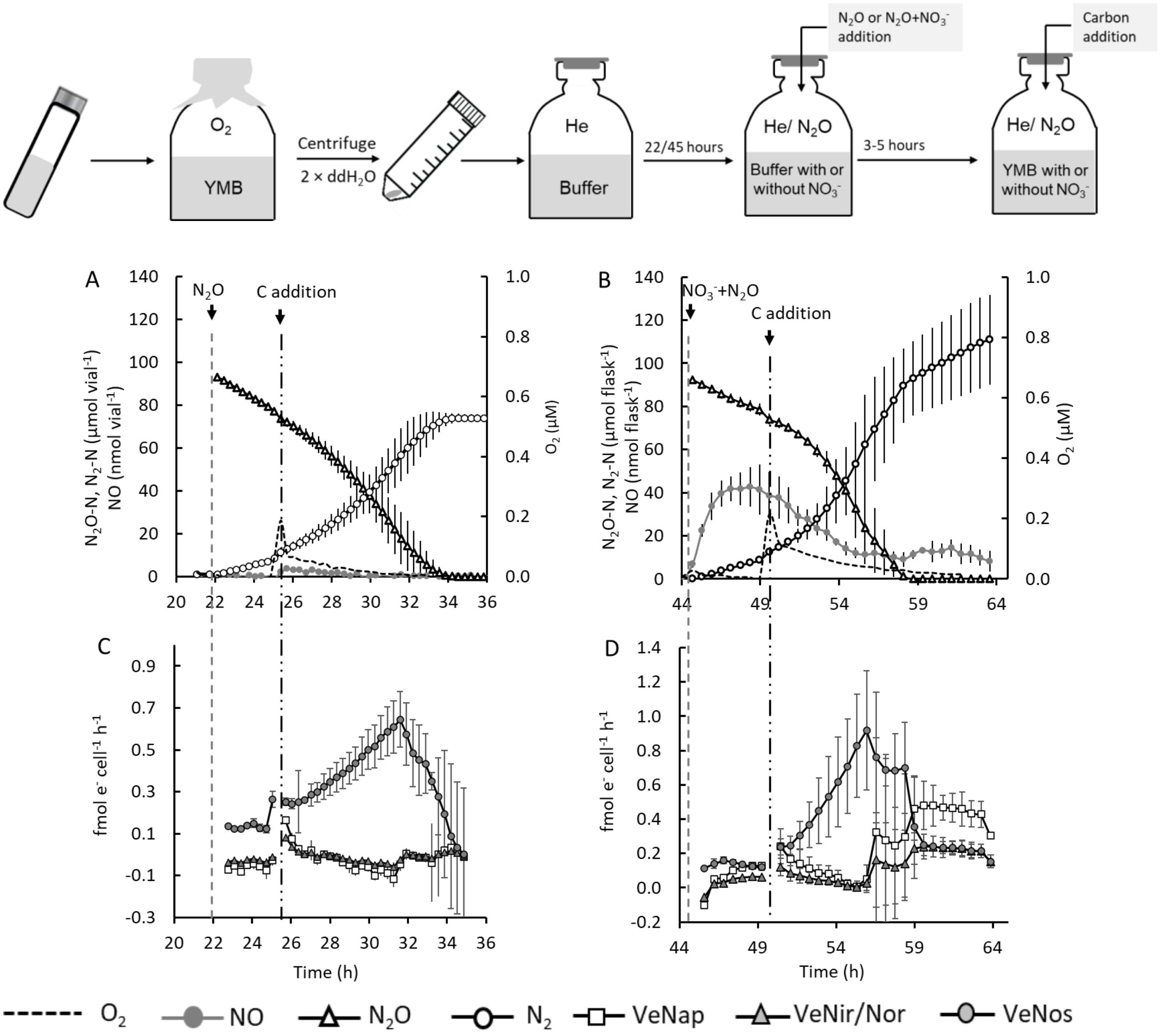


**FIG S3** Denitrification kinetics of starved cultures of *Bradyrhizobium* strain HAMBI 2125 and response to carbon addition. Gas kinetics and corresponding electron flow rates of starved cultures provided with $N_2O$ (**Panels A and C**) or $N_2O$ and $NO_3^-$ (**Panels B and D**) as electron acceptor. The bioassay protocol for preparations of the cultures, depicted above the figure panels, was similar to Bioassay 2 (Fig. 1B, main text) where the organisms had to synthesize the denitrification proteome in the absence of a carbon source. The cultures were raised from stocks in YMB medium. After five days the cultures were centrifuged and washed twice using autoclaved $ddH_2O$, after which the pellets were added into pre-prepared flasks containing C-free buffer and He in headspace. No pooling of pellets took place (as opposed to Bioassay 2). The flasks (triplicate samples for each treatment) contained 9.1-10.0E+9 cells ($N_2O$ treated cultures) or 1.50-1.52 E+10 cells (flasks with $N_2O$ + $NO_3^-$). The $O_2$ concentration was < 0.2 µM in the anoxic incubation steps having He or He and $N_2O$ in headspace. After three to five hours of incubation in buffer with N-oxides, each flask received a portion of YMB (marked with dashed-dotted line), resulting in a full-strength medium (10 g/l mannitol plus 0.5 g/l yeast extract). The negative electron flow to denitrification reductases in some sampling points may be due to minor errors in calibration of N-gas measurements, as explained in detail in the main text. Bars in all graphs show standard deviation (n=3).
